# Microbiome-related aspects of locust density-dependent phase transition

**DOI:** 10.1101/2020.09.17.301010

**Authors:** Omer Lavy, Ohad Lewin-Epstein, Yonatan Bendett, Uri Gophna, Eran Gefen, Lilach Hadany, Amir Ayali

## Abstract

Locust plagues are an ancient phenomenon, with references going back to the Old Testament. These swarming pests are notorious for their tendency to aggregate and perform long migrations, consuming vast amounts of vegetation and decimating the cultivated fields on their path. However, when population density is low, locusts will express a solitary, cryptic, non-aggregating phenotype that is not considered as an agricultural pest. Although transition of locusts from the solitary to the gregarious phase has been well studied, the shifts in the locust microbiome composition associated with this phase-transition have yet to be addressed. Here, using 16S rRNA amplicon sequencing, we compared the bacterial composition of solitary desert locusts before and after a crowding-induced phase-transition. Our findings reveal that the microbiome is altered during the phase transition. We also show that this significant change in bacterial composition includes the acquisition of a specific bacterial species - *Weissella cibaria* (Firmicutes), which has been previously shown to induce aggregation in cockroaches. Our findings led us to hypothesize that the locust microbiome may play a role in inducing aggregation behavior, contributing to the formation and maintenance of a swarm. Employing a mathematical model, we demonstrate the potential evolutionary advantage of inducing aggregation under various environmental conditions; and specifically, when the aggregation-inducing microbe exhibits a relatively high horizontal transmission rate. This is a first description of a previously unknown and important aspect of locust phase transition, demonstrating that the phase shift includes a shift in the gut and integument bacterial composition.

## Introduction

Locusts are best known for their devastating potential to cause immense damage to natural vegetation and crops. These short-horned grasshoppers (Orthoptera), belonging to the family Acrididae, frequently aggregate and migrate in swarms, decimating to crops over large areas of the developing world [1–4]. All locust species may express two very different density-dependent phenotypes (or phases): at low densities, locusts behave in a cryptic manner, actively avoiding interaction with conspecifics and generally not posing a serious threat to crops (i.e. the solitary phase). When crowded, however, locusts express the notorious gregarious phenotype, tending to aggregate into swarms, to migrate, and to consume huge amounts of vegetation, thus causing severe damage to agriculture (the gregarious phase) [1,3,5,6]. The phase transition from solitary to gregarious in the different locust species encompasses many changes, including a marked behavioral change [5–8], as well as immense physiological changes[5,6,9,10].

Among studies investigating the different characteristics of the locust density-dependent phase phenomenon, very few have addressed the microbiome-related aspects. The role of gut bacteria in maintaining swarm integrity, through the emission of locust-attracting fecal volatiles, was suggested by Dillon et al. [11,12]. Recently, Lavy et al. [13,14] reported the occurrence of phase-related different temporal dynamics in bacterial composition of the gut and the reproductive tract in the desert locust (*Schistocerca gregaria*). No study to date, however, has directly addressed the bacterial composition-related aspects of the locust density-dependent phase-transition.

Recent studies have revealed mechanisms by which the microbiome can affect host behavior [15,16], including in insects [17,18]. Other studies have explored the evolutionary perspective of microbiome-induced social interactions [19–21]. In the current controlled laboratory study, we used the desert locust (*Schistocerca gregaria*) as a model with which we directly explore the association between solitary-to-gregarious phase transition and the individual locust’s microbiome. We further constructed a mathematical model with which we examined the conditions that may favor the evolution of microbial species that induce aggregation behavior in their host. We found that microbes with an advantage in horizontal transmission would benefit from inducing increased interaction rates and aggregation in their hosts, thus improving their transmission potential. This in turn would facilitate the locust crowding behavior and, in parallel with changes in population density, could result in locust swarming.

## Methods

### Insect rearing

To obtain locusts in either the solitary or the gregarious phase, locusts were reared from up to 2h post-hatching in either solitary or crowded conditions. The gregarious and the solitary locusts were maintained in different, climate-controlled rooms, under similar ambient conditions except for density (detailed in Lavy et al.[13]). Briefly, light (10D:14L cycle) and radiant heat were supplied by electric bulbs, allowing the insect’s behavioral thermoregulation between 30 - 37°C. All locusts were fed daily with fresh wheat seedlings and dry oats.

### Experimental setup

We first sampled the gut and integument bacterial composition of mature solitary-reared males and females (t=0). We then transferred the locusts (marked individually with acrylic color on their pronotum, hind femur, and wings) to heavy crowding conditions, i.e. introducing them into a 65-liter metal cage, containing ~200 crowded-reared conspecifics locusts of similar-age, (simulating the locust transition from solitary conditions to swarm conditions; t=7). On day 7 post-transfer, we repeated the sampling of the same individuals. The crowded-reared gregarious individuals were sampled the same way and at the same time as control.

### Sampling

The abdomen of mature individuals of both phases was swabbed (Heinz Herenz, Hamburg, Germany), followed by inserting each locust, head first, into a sterile 50 ml centrifuge tube (Corning, NY, United States) for 3 h, to collect its fecal pellets. This setup prevented the insects from moving, and ensured the collection of uncontaminated feces from each individual. Swab samples and the fecal pellets were marked as “t=0” and kept in −80 C° until DNA extraction. Re-sampling at day 7 was conducted in the same manner; samples were marked “t=7” and stored at −80 C° until further use.

### DNA extraction and sequencing

Bacterial DNA was extracted using the “Powersoil” DNA isolation kit (Mo Bio Laboratories Inc., Carlsbad CA, United States) following the manufacturer’s instructions, using 60 μl for final DNA elution. To determine the bacterial composition of the different samples, a polymerase chain reaction (PCR) was applied with universal primers 5’ end common sequences (CS1-341F 5’-ACACTGACGACATGGTTCTACANNNNCCTACGGGAGGC AGCAG and CS2-806R 5’-TACGGTAGCAGAGACTTGG TCTGGACTACHVGGGTW TCTAAT), amplifying the variable regions 3 and 4 of the prokaryotic 16S rRNA gene. The PCR conditions were one cycle of 95 C° for 3 min, followed by 31 cycles of 95 C° 15 s, 55 C° 15 s, 72 C° 5 s; using the PCR master mix KAPA2G Fast™ (KAPA Biosystems, Wilmington MA, United States). Amplified products were visually verified, and sent for deep sequencing on an Illumina MiSeq platform at the Chicago Sequencing Center of the University of Illinois.

DNA was also extracted according to the protocol above from two clean swab sticks as a control. Visual validation showed no PCR product in these samples.

### Data analyses

De-multiplexed raw sequences were quality filtered (bases with a PHRED score < 20 were removed) and merged using PEAR (Zhang et al. 2014). Sequences of less than 380 bp (after merging and trimming) were discarded. Data were then analyzed using the Quantitative Insights Into Microbial Ecology (QIIME) package (Caporaso et al. 2010). Vsearch (Rognes et al. 2016) was used for chimera detection and elimination. Merged and trimmed data were additionally analyzed with the DADA2 pipeline (Callahan et al. 2016) to infer exact sequences for amplicon sequence variant (ASV) analyses, and taxonomy assignment was performed using the Silva database (version 128). To ensure data evenness, before analysis the data were rarefied to 2500 seqs/sample. All statistical analyses were conducted using “R” v.3.4.1. (R Core Team, 2013). Canonical analysis of principal coordinates and Analysis of similarities (ANOSIM) were carried out using the “vegan 2.4-3” package, (function used: “capscale”). Kruskal-Wallis rank sum test was conducted using the “stats 3.6.1” package, and Dunn’s test was conducted using the “dunn.test 1.3.5” package.

### Mathematical model description

We modeled a fully-mixed population of locusts, each bearing one of three microbiome compositions, and used a compartmental model that describes the change in the frequencies of the different microbiome compositions in the population over time. Denoted by *S* are locusts bearing a standard solitary microbiome composition. Denoted by α are locusts bearing, in addition to the standard microbiome, also microbes of type α that induce their hosts to aggregate, namely, to initiate more interactions. We denote by *g* ≥ 1 the factor that controls the fold-increase in the rate of interactions of locusts bearing microbes of type *α*. Denoted by *β* are locusts bearing, in addition to the standard microbiome, also microbes of type *β*, which do not affect their host’s behavior.

During interactions between locusts, microbes *α* and *β* can be transmitted from one locust to another with probabilities *T*_*α*_, *T*_*β*_. We assume that *α* and *β* cannot co-reside within one host, and thus upon interactions between an *α*-carrying host and a *β*-carrying host, transmission of the *α* microbes would result in replacement of the residing *β* microbes with the transmitted microbes, and vice versa for *β* to *α* transmission. We assume that the hosts bearing *α*/*β* microbes can restore their standard microbiome and lose *α*/*β* at the corresponding rates *L*_*α*_, *L*_*β*_. We focus here on the cases in which *α* microbes have a higher transmission rate than *β* microbes (*T*_*α*_ > *T*_*β*_), and a lower persistence within the microbiome (*L*_*α*_ > *L*_*β*_).

We further account for increase and decrease in *all* rates of interactions due to changes in population density, denoted by *D*. Finally, we include low rates of mutation in all directions: S ↔ *α*, S ↔ *β*, and *α* ↔ *β*. For simplicity, we assume a constant mutation rate of *μ* in all directions.

The following three differential equations represent the dynamics in the population (see Fig.3a and Supplementary Information for analysis of the system):

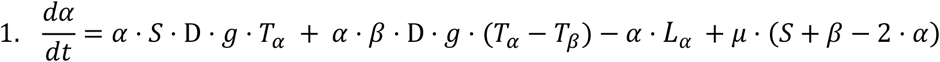

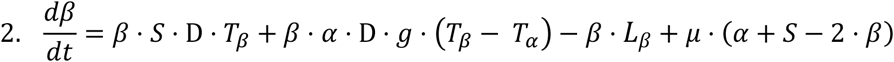

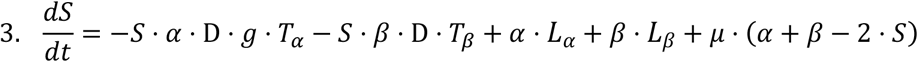

## Results

### Gregarious and solitary locust bacterial composition analysis

We successfully sequenced the integument and gut bacterial composition of 21 solitary locusts, before and after crowding them, and compared these to the integument and gut bacterial composition of 21 crowd-reared gregarious locusts. We found that the microbiome of the gregarious and solitary feces samples is comprised primarily of members of the Proteobacteria and Firmicutes (Fig.1a). The locust-integument also featured a representation of Actinobacteria and Bacteroidetes (Fig.1b).

**Figure 1.**
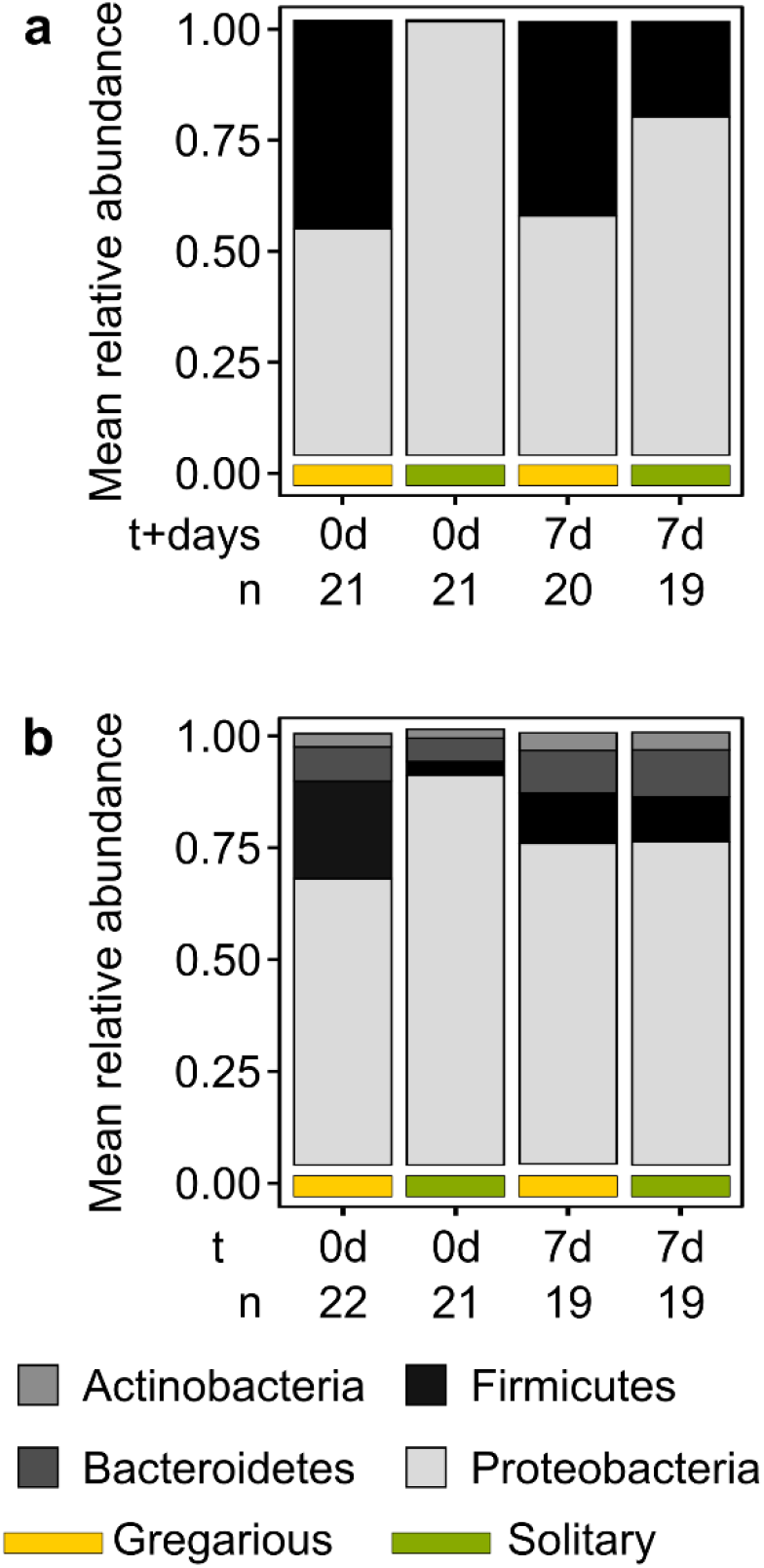
Mean relative abundance of the dominant phyla comprising the bacterial composition of the feces. **(a)** and the integument **(b)** of gregarious (yellow underscore) and solitary (green underscore) individuals at t=0d and at t=7d. It is noticeable that the phylum Firmicutes is absent from the feces of the solitary locusts at t=0 and present after seven days. In addition, while the feces bacteria of both gregarious and solitary locusts feature predominantly Firmicutes and Proteobacteria, the locust’s integument is also populated with bacteria from the Actinobacteria and Bacteriodetes phyla.

We observed that while the bacterial composition of the solitary locusts was very different from that of the gregarious control group at t=0, these differences had diminished by t=7. This noted shift in the solitary gut and integument bacterial composition could mostly be attributed to an increase in Firmicutes during the seven days of gregarious crowding (Fig. 1).

This trend of solitary-to-gregarious bacterial shift was also observed at the genus level: the integument bacterial composition of the solitary locusts notably differed from that of the gregarious locusts at t=0 days (ANOSIM: *p*<0.001, R= 0.43; Fig. 2a). By the end of the crowding period, however, the locusts of solitary origin had lost their unique bacterial composition and displayed a gregarious-like bacterial profile on their integument (ANOSIM: *p*=0.34, R= 0.005; Fig. 2a1).

**Figure 2:**
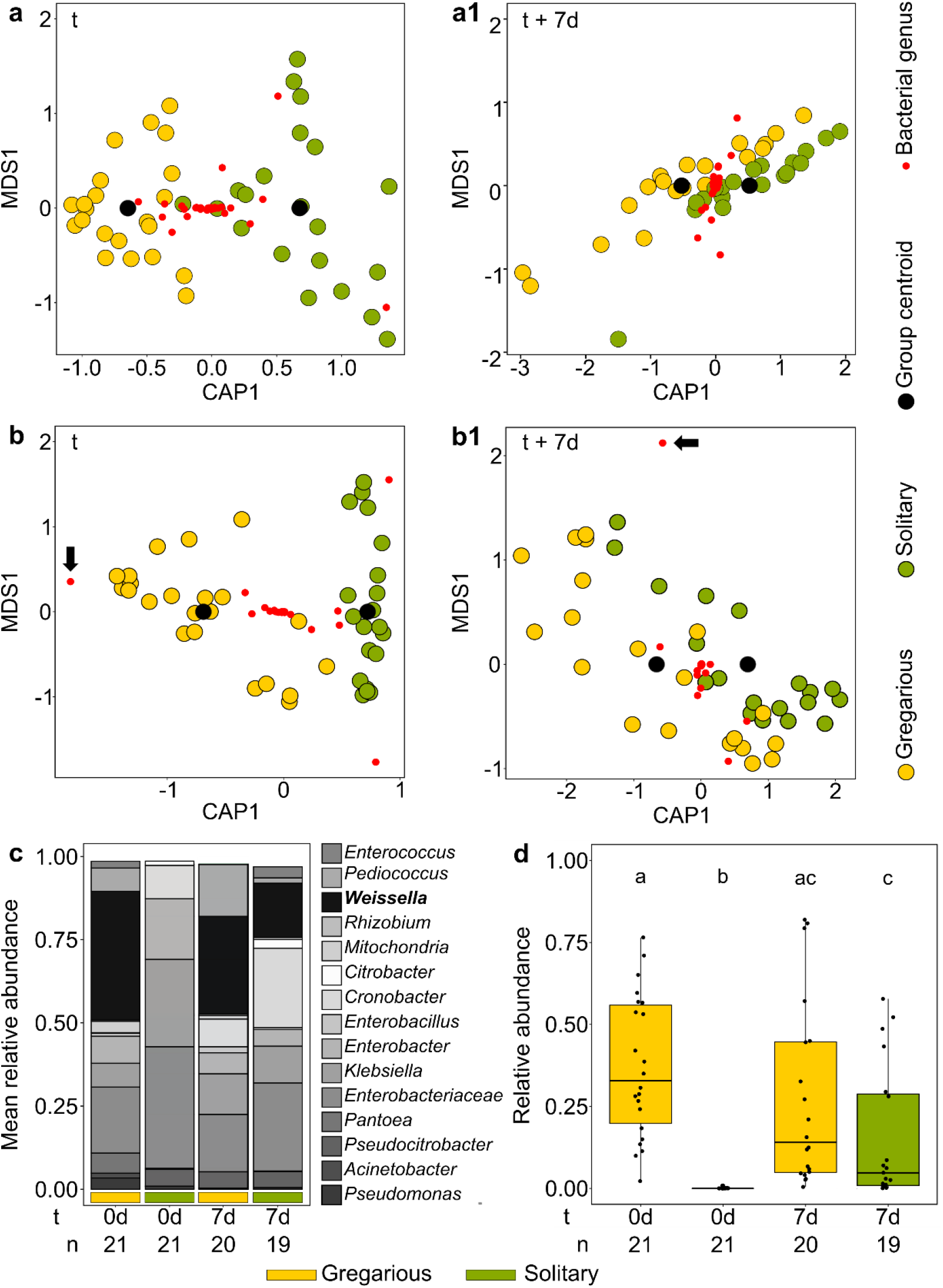
Genus level, Bray-Curtis based, Constrained Canonical Analysis of integument bacterial composition according to phase at t=0d **(a)** and t=7d **(a1)** and of the feces bacterial composition at t=0d **(b)** and at t=7d **(b1)**. The axes display the variation of the explanatory variables (i.e. the phase groups). The location of the bacterial genera (red dots) relative to the group centroids represents the between-group variability explained by a specific bacterial genus. At t=0d, the genus *Weissella* (marked with a black arrow) is a main contributor to the between-phase variance (based on its location on the X axis) in the feces bacterial composition (b); whereas, at t+7d it is located almost between the group centroids, decreasing dramatically in its contribution to the variance (b1). **(c)** Mean relative abundance of the dominant genera comprising the feces bacterial composition of gregarious (yellow underscore) and solitary (green underscore) at t=0d and at t=7d. It is noticeable that the genus *Weissella* is absent from the solitary treatment at t=0 and present after 7 days. **(d)** Relative abundance of the dominant Amplicon Sequence Variant (ASV) assigned to the genus *Weissella* in the gregarious and solitary-originated locusts. The *Weissella* assigned ASV was absent from the solitary treatment at t=0 days, whereas at t=7 days there was no significant difference between the two treatments. Differences between the groups were analyzed using Kruskal-Wallis test (*p*<0.001) and Dunn’s test as post-hoc.

A constrained canonical analysis and analysis of similarities of the locusts gut bacterial composition, revealed the same pattern of transition from a distinct solitary gut microbiota at t=0 (ANOSIM: *p*<0.001, R=0.51; Fig.2b) to bacterial compositions that were much closer between solitary and gregarious locusts after seven days of crowding (ANOSIM: *p*=0.037, R=0.08; Fig.2b1). Most notably, the genus *Weissella* (Firmicutes) was absent from the gut of solitary locusts at t=0, whereas it dominated in the gregarious population, contributing considerably to the observed per-phase difference before crowding (Fig.2b, c). In contrast, at t=7 the contribution of *Weissella* to the per-phase separation strongly decreased, as it became prevalent also among the previously solitary locusts (Fig. 2b, c).

Higher-resolution analysis of the data revealed that a specific gregarious-prevalent Amplicon Sequence Variant (ASV) had been acquired by the solitary locusts during the crowding period (Fig.2d), confirming that this is indeed the same bacterium, assigned to the species *Weissella cibaria* (top hit strain: KACC 11862, similarity: 100%), that was transmitted from the gregarious locusts to their solitary conspecifics during the gregarization process.

### Microbiome-induced aggregation model

We consider a fully-mixed population of locusts, each carrying one of three microbiome compositions: a standard solitary microbiome composition (denoted by *S*); the standard microbiome plus microbes of type *α* that induce their hosts to aggregate (denoted by *α*) - to initiate more interactions than other hosts by a factor of *g* > 1; and the standard microbiome plus microbes of type *β* which do not affect their host’s behavior (denoted by *β*). Locusts can lose microbes *α* and *β* with rates *L*_*α*_, *L*_*β*_ respectively. We use a compartmental model, as described in the *Methods* section.

We begin by analyzing the model described by equations 1-3 (see methods), constructed to examine the possibility of bacterial effects on the behavior of the locusts. We first consider populations with constant density (D(t) = 1 throughout) and without mutations (*μ* = 0). In this case we find analytical expressions for the equilibrium points of the system and their stability (see details in the Supplementary Information). The analysis revealed that in a competition between *S* and *β* alone, *β* can evolve and is maintained in polymorphism in frequency of 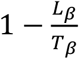 (the frequency of *S* would be 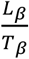). We thus see that when *T*_*β*_ ≤ *L*_*β*_, *β* goes extinct. Similarly, with α and *S*, α can evolve and be maintained in polymorphism in the frequency of 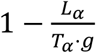 In the case of *α*, not only the horizontal transmission probability (*T*_*α*_) determines its success, but rather its effective transmission rate (*T*_*α*_ ⋅ *g*), combining both the horizontal transmission probability and the induction of interactions rate. When all three types are competing over the population, we find that *α* can be maintained in polymorphism with *S* if the following two conditions are satisfied:

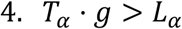

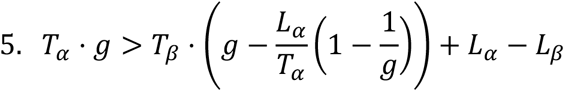

See figure 3b-c and the Supplementary Information for the derivations. We find that in the range of conditions in which α can evolve, its proportion in the population increases with the horizontal transmission advantage (*T*_*α*_/*T*_*β*_) and with the level of aggregation induced by *α* (*g*; see figure 3). This result is consistent with the empirical evidence demonstrating that the *Weissella* genus is both associated with aggregation and shows significant horizontal transmission ability.

**Figure 3.**
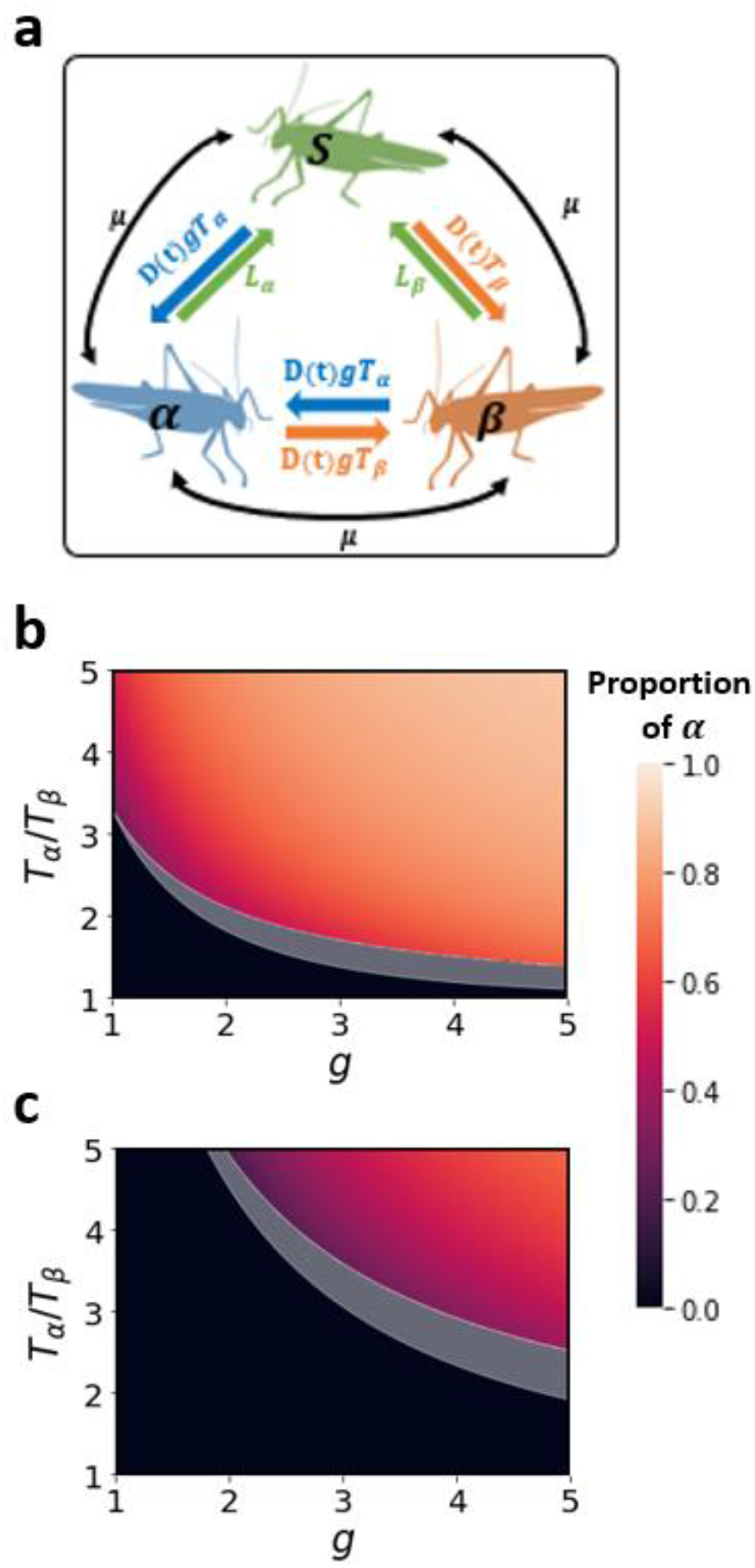
Model illustration and analytic results. **(a)** Illustration of the transition rates between the three microbiome types. *T*_*α*_ and *T*_*β*_ are the horizontal transmission rates of microbes *α* and *β*. *g* is the fold-increase in locust interactions due to the induction of aggregation generated by microbe *α*. *L*_*α*_ and *L*_*β*_ are the rates at which the locusts lose microbes *α*, *β* respectively. D(t) denotes the change in *all* interactions rate due to shifts in population density, while μ represents mutations in all possible directions. **(b-c)** The expected proportion of hosts bearing microbe α in the population, based on analysis of the equilibria of equations (1-3) and their stability (see Supplementary Information) is plotted for *L*_*α*_ = 0.06 (b) and *L*_*α*_ = 0.2 (c). The black area represents the range of parameters in which *α* cannot evolve; the grey area represents the range of parameters in which *α* will either become extinct or reach polymorphic equilibrium, depending on the initial conditions of the population; the colored area represents the range of parameters in which *α* will evolve and reach stable polymorphism with *S*, regardless of the initial conditions of the population. *T*_*β*_ = 0.025, *L*_*β*_ = 0.002, *μ* = 0, *D* = 1.

Following these results, we examined the dynamics in populations that undergo both mutations and changes in their density, e.g. due to variation in the availability of resources. We modeled a population of locusts with density D(t) that changes along time with a Gaussian wave (starting and ending at D = 1). The increase in density directly affects the rate of all interactions in the population, increasing the effective horizontal transmission rate of both microbes, *α* and *β*. We denote by *ρ* the swarming level in the population, defined as the average interaction rate, which is affected both by the change in population density (*D*) and by the fold-increase in the interaction rate of *α*-bearing hosts (*g*). We find that in this type of population dynamics, *α* can spread under a much wider range of conditions, and contribute to the swarming. However, in many cases the invasion of *α* is temporary: *α* invades the population and spreads rapidly when the density is high enough, generating a crowded swarm; and it then declines and disappears from the population as the density decreases (Fig. 4).

**Figure 4.**
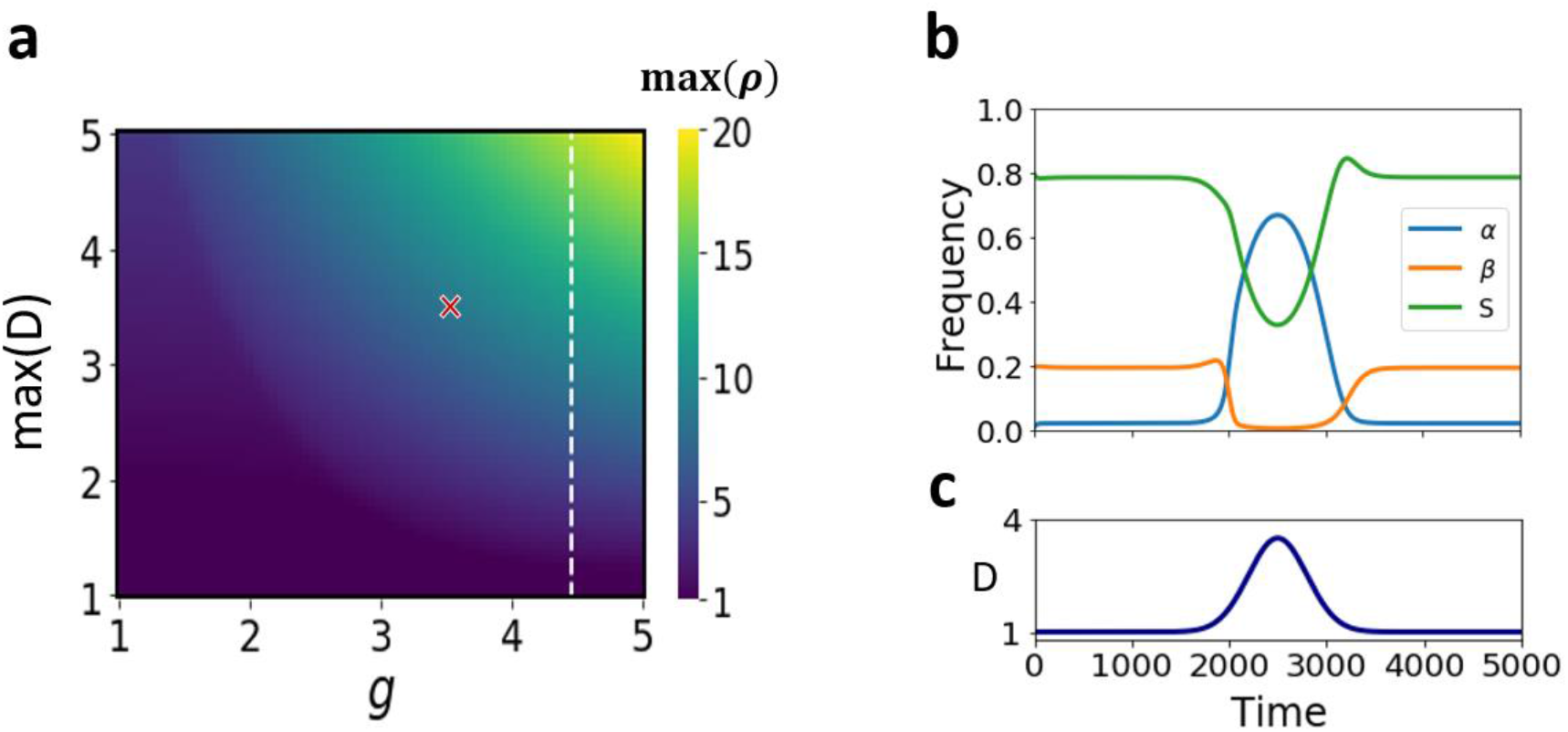
Temporary increase in the population density enables microbe α to invade and spread under a wide range of conditions. **(a)** Using numerical simulations we analyzed the swarming level in the population (ρ), defined as the mean number of interactions per locust: *ρ* = (β(*t*) + *S*(*t*)) ⋅ *D*(*t*) + α(*t*) ⋅ *g* ⋅ *D*(*t*). We calculate max *ρ* in populations that undergo a Gaussian wave of an increase followed *t* by a decrease in its density, and plot the maximal *ρ* as function of α’s induction of aggregation (*g*) and the population’s maximal density (*max*(*D*)). The area to the right of the dashed white line represents parameter values for which α reaches stable polymorphism with *S* also if *D* = 1 throughout, while the area to the left of the dashed white line represents parameter values for which α is maintained only in mutation-selection balance proportions when *D* = 1. The ‘x’ mark denotes the *max*(*D*) and *g* values used in panels (b-c). **(b)** We plot a time-series example of the results obtained in one numerical simulation run. The figure shows the proportion of the three microbiome types along time. *T*_*α*_ = 0.05, *T*_*β*_ = 0.025, *L*_*α*_ = 0.2, *L*_*β*_ = 0.02 **(c)** The change in population density (*D*) along time. Similar density functions of time were used in all simulations, with the only difference between simulations being the maximal value of *D*.

Our model thus predicts that microbes, such as *Weissella* (represented by α in our model) may “take over” the gregarious population using a combination of two effective mechanisms: the first is by enjoying a higher horizontal transmission rate than its competitors in the locust microbiome (*T*_*α*_ > *T*_*β*_); and the second is by eliciting the conspecific interactions (g) (i.e. the conspecific attraction) of infected insects; or, within the locust context, making the individuals more gregarious (Fig. 2).

## Discussion

The density-dependent phase shift of locusts from the solitary to the gregarious phenotype has been widely studied, and shown to include behavioral changes as well as physiological transitions [2,5,6,22]. However, the locust bacterial composition-related aspects of the phenomenon have never been directly investigated.

### Locust-bacteria phase shift dynamics

The bacterium *Weissella cibaria* (Firmicutes) has been shown here to dominate the gregarious locust’s bacterial composition, and to be transmitted to solitary locusts upon crowding. *Weissella* members have been previously described as dominant gut bacteria in gregarious locusts of several species, and mainly in the migratory locust (*Locusta migratoria*) [23–25] and the desert locust (*S. gregaria*) [13,26]). In Lavy et al. [13] we reported the intriguing pattern of temporal *Weissella* blooms in the gut of laboratory-reared gregarious locusts, while the solitary locusts were shown to maintain a constant bacterial composition dominated by members of the phylum Proteobacteria. The data presented here further demonstrates the ability of this genus to infect and dominate the microbiota of previously naive solitary locust-hosts when entering the gregarious phase. Hence this bacterium is suggested to be instrumental in the crowding-induced shift of the solitary locust gut bacterial composition to a gregarious-like composition, a previously unexplored aspect of locust phase-transformation.

Wada-Katsumata et al. [27] demonstrated that the presence of *W. cibaria* in cockroach feces attracts cockroach nymphs in the vicinity and elicits their aggregation. In light of those and the present findings, we speculate that *W. cibaria* may also benefit from eliciting aggregation behavior among locusts, thereby increasing locust conspecific encounters and thus also the bacterium’s own transmission rate. This is even more plausible when taking into account that, while being very common among gregarious locusts [13,23–26], this bacterium is not vertically transmitted from parent to offspring [26] and thus has to rely on horizontal transmission.

In this study we also describe for the first time the bacterial composition of the locust integument. It contains a large fraction of Proteobacteria and Firmicutes that may be partly gut-derived and a result of the dense conditions in the locust cage; we also report on a fraction of Bacteroidetes and Actinobacteria, which were not found in the fecal samples. Members of the Actinobacteria have been found to function as a first line of insect defense against pathogens.[28–30]. Consequently, we hypothesize that they may also serve the same purpose in contributing to the locust’s immunity, though this needs to be validated in further studies.

### Insect aggregation behavior can be a bacterial ecological strategy

We suggest that the microbiome may play an essential role in the locust phase-shift, and that the microbiome has the potential to promote crowding among gregarious populations. Though the controlled manipulations in a laboratory cage cannot of course fully simulate field conditions, the laboratory set-up allowed us to closely monitor individual solitary locusts following their exposure to gregarious populations and consequent phase transformation; a study which could not be achieved in the field (due to the scarce solitary individuals and migrating gregarious swarms). Furthermore, both Dillon et al. [31] and Lavy et al. [13] have shown that the bacterial composition of laboratory locusts is broadly similar to that of field-collected individuals, providing the laboratory data with further ecological relevance.

The ecological relevance of our findings is highlighted by our mathematical model. Using this model, we studied the conditions that enable the evolution of an aggregation-inducing microbe (*α*) in a locust population. We found that while benefitting from the advantage of horizontal transmission, a microbe may further benefit from inducing its host to aggregate and increase its rate of interactions, thus enabling more transmission opportunities. This induction can be beneficial even when incurring a certain cost (represented in our model by *L*_*α*_ > *L*_*β*_). We have further shown that incorporating changes in population density in the model due to external reasons, in the form of a sinuous increase followed by a decrease, allowed the aggregation-inducing microbe (*α*) to invade the population under a much wider range of conditions. This invasion was shown to be transient, characterized by a rapid spread of microbe α when the population becomes dense enough, and followed by a decline as population density decreases. The increase in α in the population in itself further increases the crowding, as *α*-bearing locusts tend to aggregate and seek additional interactions.

A solitary-to-gregarious phase-shift in the field is strongly affected by forced crowding, caused by factors such as food patchiness or a drastic population increase [5,6]. Such conditions support and induce conspecific encounters, which potentially increase the bacterial spread within the forming swarm.

As noted, Lavy et al. [13,26] indicated that although *Weissella* seems to spread very quickly in the gregarious population, it is not transmitted across generations, unlike other locust-bacteria. Our model can account for this lack of transmission across generations, if interpreted as multigenerational, considering *L*_*α*_ and *L*_*β*_, the transitions from *α* and *β* to *S*, and the lack of transitions in the opposite direction. This can represent a population in which *S* is a standard microbiome that is also transmitted across generations, while *α* and *β* cannot be transmitted or are less successful in transmitting across generations. Therefore, *α* and *β*-bearing hosts lose a certain proportion to *S* hosts (with rates *L*_*α*_ and *L*_*β*_). Such a failure to be transmitted to offspring, could be due to an inability to colonize the younger life stages of the locust, or due to lack of maternal inoculation [26].

### Concluding remarks

We have described here, using empirical experiments and a novel mathematical model, how bacteria of solitary individuals, changes upon crowding. Furthermore, we showed that some bacterial agents that can potentially benefit from the aggregation phenomenon, may play an instrumental part in the locust density-dependent phase-shift, under the controlled conditions of the laboratory. Our findings suggest that this is beneficial for the underlying bacterium, offering it a multitude of additional potential hosts. It should be noted, however, that many questions remain unanswered; and further experiments validating *Weissella*’s potential in inducing aggregation in locusts are required. This is a first description of the dynamics of the locust’s microbiota, as its host undergoes the physiological and behavioral changes associated with phase-transition; and our findings illuminate previously unexplored aspects of this quintessential example of environmentally-induced plasticity. Our findings may also open the way to potentially novel directions in the on-going efforts to battle this ancient and devastating pest.

