## Supplementary Information for "Microbiome-related aspects of locust density-dependent phase transition"

### Supplementary Note 1 The microbiome-induced aggregation model

We use a compartmental model to describe the change in the frequencies of three different microbiome compositions in a fully-mixed locust population over time.

Hereby are the model parameters:

| | |
|---|---|
| $S$ | Proportion of hosts carrying a standard microbiome |
| $\alpha$ | Proportion of hosts carrying a standard microbiome plus microbes of type $\alpha$ |
| $\beta$ | Proportion of hosts carrying a standard microbiome plus microbes of type $\beta$ |
| $g$ | The fold-increase in interaction rate of hosts carrying microbes of type $\alpha$ |
| $T_\alpha$ | Transmission and establishment probability of microbe $\alpha$ during hosts interaction |
| $T_\beta$ | Transmission and establishment probability of microbe $\beta$ during hosts interaction |
| $L_\alpha$ | The rate at which hosts carrying microbes of type $\alpha$ lose the microbe, and return to carrying the standard microbiome alone. |
| $L_\beta$ | The rate at which hosts carrying microbes of type $\beta$ lose the microbe, and return to carrying the standard microbiome alone. |
| $D$ | The fold-increase in interaction rate of all hosts due to environmental factors |
| $\mu$ | Mutation rate in all directions: $S \leftrightarrow \alpha$ , $S \leftrightarrow \beta$, and $\alpha \leftrightarrow \beta$ |

Following the model settings, described in the **Methods** section of the main text, we derive three differential equations that represent the dynamics in the population.

$$S1.\quad \frac{d\alpha}{dt} = f_1(\alpha,\beta,S) = \alpha \cdot S \cdot \mathrm{D} \cdot g \cdot T_\alpha \;+\; \alpha \cdot \beta \cdot \mathrm{D} \cdot g \cdot (T_\alpha - T_\beta) - \alpha \cdot L_\alpha \;+ \mu \cdot (S + \beta - 2 \cdot \alpha)$$

$$S2.\quad \frac{d\beta}{dt} = f_2(\alpha,\beta,S) = \beta \cdot S \cdot \mathrm{D} \cdot T_\beta + \beta \cdot \alpha \cdot \mathrm{D} \cdot g \cdot \left(T_\beta - T_\alpha\right) - \beta \cdot L_\beta + \mu \cdot (\alpha + S - 2 \cdot \beta)$$

$$S3.\quad \frac{dS}{dt} = f_3(\alpha,\beta,S) = -S \cdot \alpha \cdot \mathrm{D} \cdot g \cdot T_\alpha - S \cdot \beta \cdot \mathrm{D} \cdot T_\beta + \alpha \cdot L_\alpha + \beta \cdot L_\beta + \mu \cdot (\alpha + \beta - 2 \cdot S)$$

while $\alpha + \beta + S = 1$

### Supplementary Note 2 Equilibria analysis

For the equilibrium analysis, we focus on populations with constant density $(D(t) = 1)$ and without mutations $(\mu = 0)$. Thus, equations S1-S3 are simplified to:

$$S4.\ \ \frac{d\alpha}{dt} = f_1(\alpha, \beta, S) = \alpha \cdot S \cdot g \cdot T_\alpha \ + \ \alpha \cdot \beta \cdot g \cdot (T_\alpha - T_\beta) - \alpha \cdot L_\alpha$$

$$S5.\ \ \frac{d\beta}{dt} = f_2(\alpha, \beta, S) = \beta \cdot S \cdot T_\beta + \beta \cdot \alpha \cdot g \cdot \left(T_\beta - \ T_\alpha\right) - \beta \cdot L_\beta$$

$$S6.\ \ \frac{dS}{dt} = f_3(\alpha, \beta, S) = -S \cdot \alpha \cdot g \cdot T_\alpha - S \cdot \beta \cdot T_\beta + \alpha \cdot L_\alpha + \beta \cdot L_\beta$$

We find that the system contains the following four equilibria:

$$S7.\ \ (\alpha, \beta, S) = (0,0,1)$$

$$S8.\ \ (\alpha, \beta, S) = \left(1 - \frac{L_\alpha}{T_\alpha \cdot g}\ , 0\ ,\ \frac{L_\alpha}{T_\alpha \cdot g}\right)$$

$$S9.\ \ (\alpha, \beta, S) = \left(0, 1 - \frac{L_\beta}{T_\beta}, \frac{L_\beta}{T_\beta}\right)$$

$$S10.\ \ (\alpha, \beta, S) = \left(\begin{array}{c} \dfrac{L_\alpha - L_\beta \cdot g - T_\alpha \cdot g + T_\beta \cdot g}{(T_\alpha - T_\beta) \cdot g \cdot (g-1)}, \\ \dfrac{-L_\alpha \cdot T_\alpha \cdot g + L_\alpha \cdot T_\beta \cdot g - L_\alpha \cdot T_\beta + L_\beta \cdot T_\alpha \cdot g + T_\alpha^2 \cdot g^2 - T_\alpha \cdot T_\beta \cdot g^2}{T_\beta \cdot g \cdot (T_\alpha - T_\beta) \cdot (g-1)}, \\ \dfrac{L_\alpha - L_\beta - T_\alpha \cdot g + T_\beta \cdot g}{T_\beta \cdot (g-1)} \end{array}\right)$$

While equilibrium (S7) is trivial, representing fixation of $S$, equilibria S8 and S9 are polymorphic, maintaining $S$ alongside with either $\alpha$ (S8) or $\beta$ (S9). We can see that S8 is valid (containing three non-negative values that sum up to 1) only if:

$$S11.\ \ T_\alpha \cdot g > L_\alpha$$

and S9 is valid only if:

$$S12.\quad T_\beta > L_\beta$$

The fourth equilibrium (S10) is also polymorphic and includes all three microbiome types, nevertheless it is not valid for the parameter range we focus on, where $g > 1$ and $T_\alpha > T_\beta$ (numerically validated over more than $10^9$ parameter sets in the range of $0 < T_\alpha, T_\beta, L_\alpha, L_\beta < 1$ and $1 < g < 20$).

### Supplementary Note 3 Equilibria stability analysis

To analyze the stability of the equilibria, we derive the Jacobian matrix of the system $(f_1, f_2, f_3)$ with respect to the variables $\alpha, \beta, S$ and determine the eigenvalues of the Jacobian in each equilibrium point. We find that the Jacobian matrix is:

$$S13.\quad J = \begin{pmatrix} \frac{\partial f_1}{\partial \alpha} & \frac{\partial f_1}{\partial \beta} & \frac{\partial f_1}{\partial S} \\ \frac{\partial f_2}{\partial \alpha} & \frac{\partial f_2}{\partial \beta} & \frac{\partial f_2}{\partial S} \\ \frac{\partial f_3}{\partial \alpha} & \frac{\partial f_3}{\partial \beta} & \frac{\partial f_3}{\partial S} \end{pmatrix}$$

$$= \begin{pmatrix} SgT_\alpha + \beta g(T_\alpha - T_\beta) - L_\alpha & \alpha g(T_\alpha - T_\beta) & \alpha g T_\alpha \\ \beta g(T_\beta - T_\alpha) & ST_\beta + \alpha g(T_\beta - T_\alpha) - L_\beta & \beta T_\beta \\ L_\alpha - SgT_\alpha & L_\beta - ST_\beta & -\alpha g T_\alpha - \beta T_\beta \end{pmatrix}$$

The eigenvalues of $J|_{S7}$ (the Jacobian at equilibrium S7: fixation of $S$) are all negative, and thus equilibrium S7 is stable, if:

$$S14.\quad T_\alpha g < L_\alpha \text{ and } T_\beta < L_\beta$$

The eigenvalues of $J|_{S8}$ (the Jacobian at equilibrium S8: polymorphism of $\alpha$ and $S$) are all negative, and thus equilibrium S8 is stable if:

$$S15. \;\; T_\alpha g > L_\alpha \;\; \text{and} \;\; \frac{L_\alpha T_\beta}{T_\alpha} + L_\beta + T_\alpha g > \frac{L_\alpha T_\beta}{T_\alpha g} + L_\alpha + T_\beta g$$

The eigenvalues of $J|_{S9}$ (the Jacobian at equilibrium S9: polymorphism of $\beta$ and $S$) are all negative, and thus equilibrium S9 is stable if:

$$S16. \;\; T_\beta > L_\beta \;\; \text{and} \;\; L_\alpha + T_\beta g > T_\alpha g + L_\beta g$$

**We thus find that $\alpha$ can be maintained in polymorphism if S15 is satisfied (conditions (4) and (5) in the main text), but it can also go extinct if S16 is maintained, depending on the initial conditions of the population. If S15 is maintained and S16 is not maintained, the extinction of $\alpha$ is not stable and thus $\alpha$ is expected to evolve and reach polymorphism.**
